# Phytosulfokine signaling modulates salt stress responses and cell wall remodeling in Arabidopsis

**DOI:** 10.64898/2026.09.07.749821

**Authors:** Joy Debnath, Zhang Jiang, Tom van der Meer, Mark Roosjen, Nadja Braun, Dolf Weijers, Christa Testerink, Timo Engelsdorf, Nora Gigli Bisceglia

**Affiliations:** Plant Stress Resilience, Institute of Environmental Biology, Utrecht University, Utrecht, the Netherlands; Laboratory of Plant Physiology, Wageningen University & Research, Wageningen, the Netherlands; Biochemistry, Wageningen University, Wageningen University & Research, the Netherlands; Molecular Plant Physiology, Department of Biology, Marburg University, 35043 Marburg, Germany

**Keywords:** Apoplastic proteomics, salinity stress, cell wall integrity, cell wall remodeling, subtilisin-like proteases, phytosulfokine, signaling peptides

## Abstract

Salinity severely impairs plant growth and development. Increasing evidence suggests that the cell wall (CW) plays a central role in salt stress acclimation, not only as a structural barrier but also as a dynamic sensor that activates downstream stress signaling pathways. To identify extracellular factors involved in cell wall remodeling and integrity signaling during salt stress, we performed a comparative proteomic analysis of the Arabidopsis seedling apoplast. Among the proteins that accumulated under salt stress, we focused on three members of the Subtilisin-Like Protease family (SBT1.1, SBT1.6, and SBT5.3), which have previously been implicated in the processing of signaling peptides and CW-associated proteins. Functional analyses revealed that *sbt1.1* and *sbt5.3* mutants exhibit enhanced lignin deposition under salt stress, suggesting altered CW remodeling during stress application. Given the established role of SBT1.1 in processing PHYTOSULFOKINE (PSK) peptides, we investigated the involvement of PSK signaling in salt stress responses. Mutants lacking the two PSK RECEPTORS (*pskr1 pskr2*) displayed reduced growth and increased lignification under salt treatment. Exogenous application of PSK attenuated salt-induced responses, including MITOGEN ACTIVATED PROTEIN KINASE 6 (MPK6) phosphorylation, salt stress marker gene expression, and lignin accumulation, ultimately promoting root elongation under salt stress. Furthermore, PSK treatment mitigated salt-induced changes in CW composition. In cell wall integrity (CWI) mutants, PSK treatment failed to restore wild-type root growth under salt stress and induced a pronounced root bending phenotype in *fer-4*, indicating that CWI signaling influences PSK-mediated root growth responses. Together, these results support a role for PSK signaling in modulating CW-associated responses to salinity stress in Arabidopsis.

## Introduction

The plant cell wall (CW) represents a key cellular component required to coordinate responses to both biotic and abiotic stress (Debnath et al., 2026; Le Gall et al., 2015; Tenhaken, 2015). This complex extracellular matrix is highly dynamic and essential for ensuring structural integrity while undergoing extensive remodeling to maintain its integrity under stress conditions (Debnath et al., 2026). CW constituents are complex polysaccharides organized into three major classes: cellulose, hemicellulose, and pectin (Delmer et al., 2024). In combination with structural proteins and aromatic polymers (lignin and suberin), these components form a highly dynamic and interconnected network capable of sustaining diverse mechanical properties across developmental stages and in response to environmental conditions (Gibson, 2012). In dicotyledonous species, the primary CW contains relatively low levels of cellulose (typically below 20%) (Gigli-Bisceglia et al., 2020). The stiffness of cellulose arises from its highly ordered crystalline structure (Arockiasamy et al., 2025). Cellulose consists of parallel β-1,4-D-glucan chains synthesized at the plasma membrane by Cellulose Synthases (CESAs) (Desprez et al., 2007; Persson et al., 2007; Taylor et al., 2000).

In *Arabidopsis*, cellulose microfibrils are associated with pectin, the most abundant component of primary CW in dicots, which plays a central role in regulating cell expansion and CW elasticity (Cosgrove, 2022). Homogalacturonan (HG), a linear polymer composed of α-1,4-linked D-galacturonic acid residues, constitutes the major structural domain of pectin (Anderson and Pelloux, 2025; Atmodjo et al., 2013; Ridley et al., 2001). The degree of HG methylesterification (DM) is dynamically regulated within the cell wall through the activity of Pectin Methylesterases (PMEs) and their Inhibitors (PMEIs) (Gallemí et al., 2022; Wormit and Usadel, 2018). Reduced methylesterification, mediated by PMEs, can promote Ca²⁺-HG-dependent cross-links, leading to increased wall rigidity through the formation of so-called “egg-box” structures (Morris et al., 1982; Obomighie et al., 2025; Zdunek et al., 2021). Recent studies highlight alternative cross-linking arrangements and emphasize the importance of pectin organization and spatial distribution in determining wall extensibility (Haas et al., 2021, 2020; Obomighie et al., 2025). Indeed, depending on the biochemical and mechanical context, increased PME activity can promote either wall stiffening or loosening (Debnath et al., 2026; Peaucelle et al., 2011, 2008; Wang et al., 2020). The absence of a strict linear correlation between PME activity and cell wall stiffness reflects the involvement of additional processes, including homogalacturonan degradation mediated by Polygalacturonases (PGs) and Pectate Lyases (PLs), as well as HG-HG cross-linking facilitated by the availability of apoplastic Ca²⁺ ions; both processes are influenced by PME activity (Huang et al., 2022; Tibbits et al., 1998; Wakabayashi et al., 2003). In addition to HG, pectins include two structurally complex branched domains: rhamnogalacturonan-I (RG-I) and rhamnogalacturonan-II (RG-II) (L. Zhang et al., 2025). The latter contains a homogalacturonan backbone substituted with the sugar apiose, which can form dimers through borate diester cross-links, influencing cell wall mechanical properties (Delmer et al., 2024; Hays et al., 2025; O’Neill et al., 2004).

Plants sense and respond to alterations in CW composition through a mechanism commonly referred to as the cell wall integrity (CWI) maintenance mechanism (Gigli-Bisceglia et al., 2020; Vaahtera et al., 2019). This mechanism has been extensively studied following inhibition of CESAs by the cellulose biosynthesis inhibitor Isoxaben (ISX) (Amanda and Hamann, 2023; Bacete et al., 2022; Denness et al., 2011; Engelsdorf et al., 2018; Gigli-Bisceglia et al., 2018; Paredez et al., 2008; Zhai et al., 2024). Plant responses to ISX require a combination of membrane-localized receptors and channels capable of triggering intracellular signaling cascades, likely linked to mechanical distortions caused by changes in CW mechanics resulting from cellulose impairment (Baez et al., 2022; Vaahtera et al., 2019; Wolf, 2022). Most CWI receptors described to date belong to the *Catharanthus roseus* Receptor-Like Kinase1-Like (*Cr*RLK1L) family, which comprises 17 members in *Arabidopsis*. Among these, FERONIA (FER), THESEUS1 (THE1), and HERCULES RECEPTOR KINASE 1 (HERK1) have been implicated in cell wall damage signaling, with THE1 being the main regulator of downstream responses to ISX-triggered cellulose biosynthesis inhibition (Bacete and Hamann, 2020; Engelsdorf et al., 2018; Feng et al., 2018; Gigli-Bisceglia et al., 2022, 2018; Richter et al., 2017; Wolf, 2022; Zhai et al., 2024). These receptors contain extracellular malectin-like domains capable of interacting with both peptide and cell wall ligands (Ge et al., 2019; Nissen et al., 2016). FER, for example, binds de-methylesterified pectin, similarly to BUDDHA’S PAPER SEAL 1 (BUPS1) and ANXUR1/2 (ANX1/2) (Feng et al., 2018; Lin et al., 2022). *Cr*RLK1Ls also act as receptors for the peptides Rapid Alkalinization Factors (RALFs), which function as signaling molecules as well as structural components of the cell wall and, in some cases, are associated with de-methylesterified pectin, likely due to their chemical properties and charge (Baillie et al., 2024; Blackburn et al., 2020; Ge et al., 2019, 2017; Moussu et al., 2023; Schoenaers et al., 2024). CW-localized Leucine-Rich Repeat Extensins (LRXs), are also able to bind the CW and interact with *Cr*RLK1Ls and RALFs (Mecchia et al., 2017; Moussu et al., 2020; Zhao et al., 2021). Salinity induces extensive changes in CW composition and organization, including alterations in pectin methylesterification and extensin arabinosylation, which affect wall porosity and activate intracellular signaling pathways (Feng et al., 2018; Gigli-Bisceglia et al., 2022; van Zelm et al., 2020; Zou et al., 2024). In halophytes such as *Salicornia*, salinity-induced CW softening increases wall elasticity and promotes cell expansion, thereby facilitating vacuolar Na⁺ sequestration and contributing to salt tolerance (Cárdenas Pérez et al., 2024).

In Arabidopsis, the *Cr*RLK1L receptor FERONIA (FER) plays a central role in coordinating responses to salt-induced CW perturbations. *fer-4* mutants exhibit severe salt hypersensitivity characterized by enhanced Mitogen Activated Protein Kinase 6 (MPK6) phosphorylation, increased expression of salt-responsive genes, and root cell bursting (Feng et al., 2018; Gigli-Bisceglia et al., 2022). Notably, these phenotypes can be alleviated by treatments that strengthen pectin networks through homogalacturonan and rhamnogalacturonan-II cross-linking, suggesting that FER monitors or stabilizes pectin-dependent wall properties under saline conditions (Feng et al., 2018; Gigli-Bisceglia et al., 2022). Similarly, mutations affecting the CW-associated proteins LRX3, LRX4, and LRX5 result in salt hypersensitivity, further supporting a role for extracellular CW signaling during salinity stress (Zhao et al., 2018).

Peptide-mediated signaling has emerged as an important component of these responses. Members of the RALF family interact with *Cr*RLK1L receptors and are closely linked to CW regulation. Salt stress promotes processing of the inactive RALF22 precursor by the SUBTILISIN-LIKE PROTEASE (SBT)6.1/S1P, generating an active peptide involved in downstream signaling (Zhao et al., 2018). In addition, RALF peptides interact with de-methylesterified pectin and influence pectin methylesterase activity, further linking peptide perception to CW remodeling (Biermann et al., 2025; Moussu et al., 2023; Rößling et al., 2024; Schoenaers et al., 2024). Other peptide families, including SERINE-RICH ENDOGENOUS PEPTIDES (SCOOPs) functioning through the Leucin-Rich-Repeat Receptor-Like Kinase (LRR-RLK) MALE DISCOVERER 1-INTERACTING RECEPTOR-LIKE KINASE 2 (MIK2) and PLANT ELICITOR PEPTIDES (PEPs), recognized by the PEP RECEPTORS (PEPR1 and PEPR2), regulate processes such as immunity, development, and CW damage responses (Bartels and Boller, 2015; Engelsdorf et al., 2018; Hou et al., 2021; Morton et al., 2025; Rhodes et al., 2021; Zhai et al., 2024). Salt stress induces the expression of PEP precursors (*ProPEPs*), and exogenous application of PEP3 can alleviate salt-induced growth defects in a PEPR1/2-dependent manner (Gigli-Bisceglia et al., 2022; Nakaminami et al., 2018; Saijo et al., 2021). Similar to RALFs, SCOOPs are synthesized as inactive precursors that require proteolytic cleavage to become active (Debnath et al., 2026). Members of the SBT3 family are required for SCOOP maturation, and loss of their activity reduces active peptide accumulation, impairing MIK2-dependent salt stress responses (Yang et al., 2023). In addition to peptide processing, SBTs also target other substrates, including PMEs (Coculo et al., 2023; Sénéchal et al., 2014). Recent work further suggests convergence between peptide signaling and CWI pathways. ISX treatment induces *SCOOP* precursor peptides transcription, where SCOOP18 contributes to CW damage signaling in a THE1-dependent manner (Zhai et al., 2024). Although mutation of THE1 alone does not alter responses to salt stress (Van der Does et al., 2017), combining the hypermorphic allele *the1-4* with loss-of-function of HERK1 (*herk1*) increases salt sensitivity (Gigli-Bisceglia et al., 2022). How this mechanism is regulated remains unclear, suggesting that *Cr*RLK1L members may function cooperatively, possibly through receptor complex formation or stabilization. These observations further support the idea that CW modifications induced in response to salt stress contribute to the activation of downstream responses. In fact, increasing CW strength, for example by enhancing HG-HG cross-links through calcium ions or by promoting borate diester bonds in RG-II, can alleviate salt stress effects (Feng et al., 2018; Gigli-Bisceglia et al., 2022).

Despite growing evidence linking CW remodeling and peptide signaling to salinity adaptation, the molecular events occurring in the apoplast during salt stress remain poorly understood. To identify apoplastic regulators of salt responses, we performed a comparative proteomic analysis of the apoplast of Arabidopsis seedlings exposed to NaCl. Among the proteins that accumulated under salt stress, we focused on three members of the SBT family (Schaller et al., 2018), SBT1.1, SBT1.6, and SBT5.3. SBT1.1 is known to process PHYTOSULFOKINE 4 (PSK4), a precursor of the sulfated peptide PHYTOSULFOKINE (PSK), which regulates plant growth, development, and stress responses (Li et al., 2024; Matsubayashi and Sakagami, 1996; Mosher et al., 2013; Sauter, 2015; Stührwohldt et al., 2021). Six PSK precursor genes (*ProPSK1-6*) have been identified, all sharing a highly conserved pentapeptide sequence, Y(SO₃H)-I-Y(SO₃H)-T-Q (Kaufmann et al., 2021; Kaufmann and Sauter, 2019; Kutschmar et al., 2009). Tyrosylprotein sulfotransferases (TPSTs) catalyze the sulfation of different PSK variants (-α, -γ, -δ, and -ε), among which PSK-α is widely found in seed plants (Z. Zhang et al., 2025). Notably, PSK-α has been shown to influence CW reconstruction during protoplast regeneration in *Daucus* species (Godel-Jędrychowska et al., 2019). PSK signaling is perceived by the plasma membrane-localized Leucine-Rich Repeat Receptor-Like Kinases PSKR1 and PSKR2 (Matsubayashi et al., 2006; Z. Zhang et al., 2025). To investigate the role of phytosulfokine in salt stress and its potential link to SBT1.1 abundance upon salt, we characterized salt responses in the double mutant *pskr1-3 pskr2-1* (Stührwohldt et al., 2021). We found that *pskr1-3 pskr2-1* seedlings display impaired growth, reduced expression of salt-responsive genes, and increased lignin deposition upon salt treatment. In contrast, PSK application reduced NaCl-induced MPK6 phosphorylation, decreased stress-related gene expression, and alleviated salt-induced changes in CW composition, including lignin accumulation, ultimately improving growth under saline conditions.

## Results

### Salt stress alters the apoplastic proteome of Arabidopsis seedlings

To identify proteins involved in salt stress responses, we performed apoplastic proteomics on six-day-old Col-0 (wild type) *Arabidopsis* seedlings treated for 24 hours with or without 100 mM NaCl in liquid medium (Figure 1A). Since salt stress may compromise plasma membrane and CWI, we also analyzed proteins accumulating in the treatment medium to characterize the extracellular protein fraction under mock and salt stress conditions (Figure 1A/B, S1; Dataset S1; Table S1). For both apoplastic extracts and medium-derived proteins, enrichment was performed using filter aided sample preparation (FASP) (Wiśniewski et al., 2009). We generated a Venn diagram including only proteins that passed the 0.05 FDR threshold (Figure 1B, 1C, Table S1). Although proteins detected in the growth medium may include biologically relevant extracellular proteins (35) (Figure S1, Table S1/S2, Dataset S1), we focused on proteins retained in the apoplast (74) (Figure 1D, Table S2), which are more likely to represent regulated extracellular responses and less likely to be caused by stress-induced leakage. GO Cellular Component analysis confirmed that most proteins identified were annotated to the apoplast, while a minority was assigned to intracellular compartments, including the nucleolus, cytoplasmic stress granules, and plastid- and ER-related compartments (Figure 1E). Within the NaCl vs. mock apoplast contrast list, we identified several putative CW-associated enzymes and Subtilisin-Like Serine Proteases (SBTs). We focused on three SBTs (AT4G34980, AT1G01900, and AT2G04160), all of which accumulated significantly in the apoplast following salt treatment (Figure 1D). To determine whether changes in protein abundance were associated with transcriptional regulation, we performed a salt stress time-course analysis of candidate gene transcript amounts (Figure S2A). Among the selected candidates, only *SBT1.1* and *SBT5.3* showed a statistically significant increase in gene expression at 3 h under salt stress compared to their corresponding controls. In contrast, *SBT1.6* transcript levels remained unchanged, indicating that its increased abundance in the apoplast is unlikely to result from transcriptional regulation. SBT1.1 has been shown to process the phytosulfokine precursor PSK4 (Srivastava et al., 2008). We therefore investigated whether salt stress affects the expression of *PSK* precursor transcripts (*ProPSKs; ProPSK1-6*) and their receptors, *PSKR1* and *PSKR2* (Figure S2B/C). We observed that *ProPSK4* is induced during early salt stress but becomes strongly downregulated over time. Other PSK precursors also responded to salt stress. For example, *ProPSK1* and *ProPSK3* were induced at 3 h, while *ProPSK5* showed early accumulation at 1 h. In addition, the transcripts of the receptors *PSKR1* and *PSKR2* accumulated at 3 h (Figure S2C).

**Figure 1.**
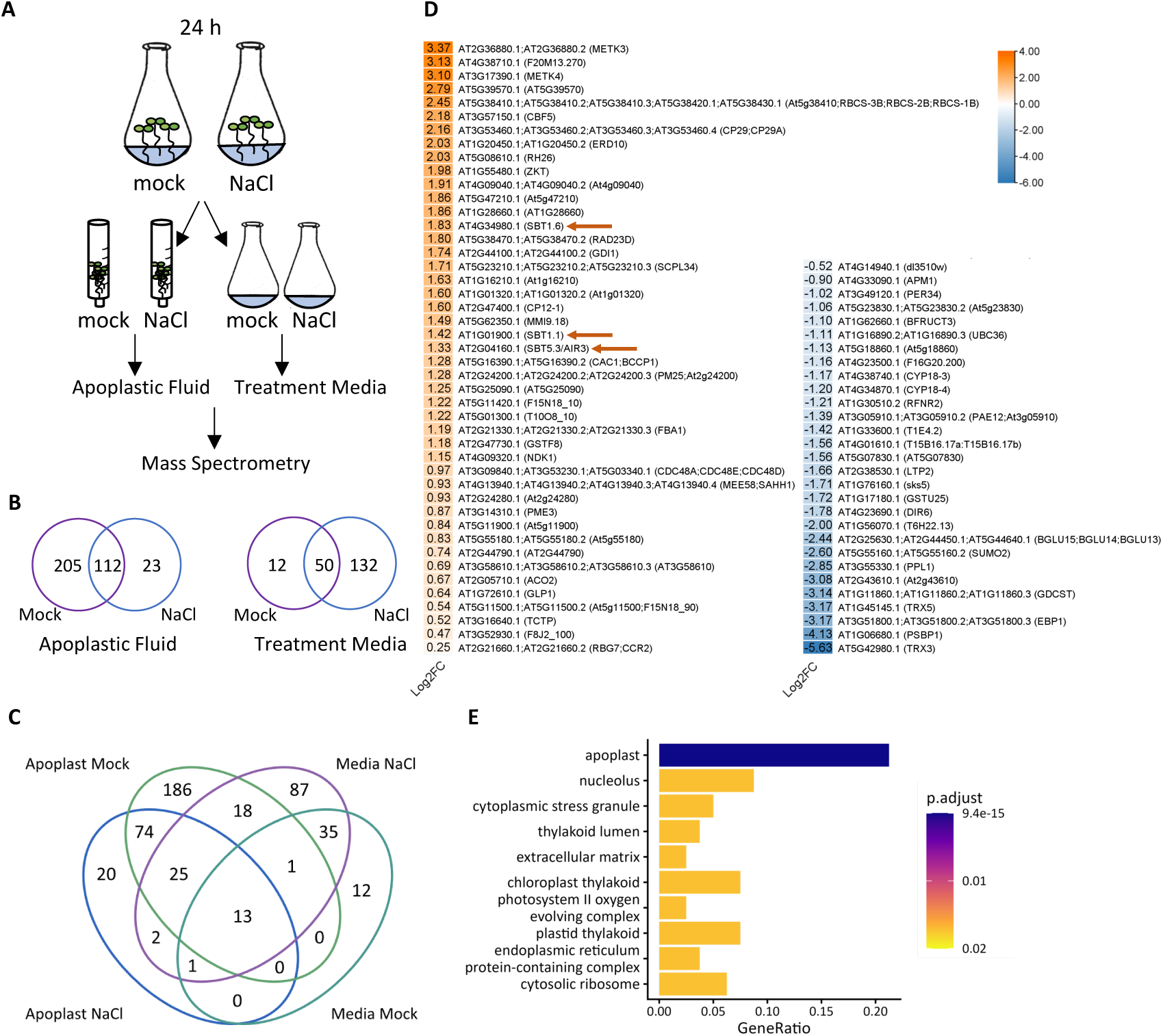
Apoplastic proteomic analysis of Arabidopsis seedlings reveals differentially abundant proteins under salt stress. (A) Schematic overview of the experimental design. Six-day-old Col-0 seedlings were treated for 24 h with either MS/2 medium (mock) or MS/2 medium supplemented with 100 mM NaCl. Three independent biological replicates were analyzed per treatment, each consisting of approximately 1 g of seedlings (n = 3). Apoplastic and medium-leaked proteins were collected, enriched using 10-kDa cutoff filters, and identified by mass spectrometry. (B) Differentially accumulated proteins (FDR ≤ 0.05) were grouped according to two comparisons: NaCl vs. mock in the apoplast fraction and NaCl vs. mock in the medium fraction. The number of proteins identified in each comparison and their overlap are shown in the Venn diagram. (C) Four-set Venn diagram showing the overlap among proteins identified in the apoplast and medium fractions under mock and NaCl conditions. (D) Heatmap showing the log₂ fold change of 74 apoplastic proteins differentially accumulated between mock- and NaCl-treated seedlings. Venn diagrams and the heatmap were generated using TBtools v2.152 (Chen et al., 2020). Arrows indicate candidate proteins selected for further analysis in this study. (E) Gene Ontology (GO) Cellular Component (CC) enrichment analysis of the proteins shown in (D), performed using the clusterProfiler package and visualized using enrichplot in R (Wu et al., 2021; Yu et al., 2012).

### SBT1.1 and SBT5.3 contribute to early salt stress signaling and regulate salt-induced lignification

To functionally characterize the role of the selected candidate genes in salt stress responses, we characterized several T-DNA insertion lines (Figure S3, Table S3). We first examined whether these candidates contribute to early salt stress signaling by analyzing the expression of three marker genes in 6-day-old seedlings treated with 100 mM NaCl for 1 h (Figure 2A). These included *REDOX RESPONSIVE TRANSCRIPTION FACTOR1 (RRTF1)*, a salt-responsive marker (Gigli-Bisceglia et al., 2022; Lamers et al., 2025; Soliman and Meyer, 2019), *ProPEP3*, associated with CW damage responses (Engelsdorf et al., 2018; Gigli-Bisceglia et al., 2022, 2018), and *WRKY DNA-BINDING PROTEIN 40* (*WRKY40*), which is involved in general stress and ABA signaling (Geilen and Böhmer, 2015; Gigli-Bisceglia et al., 2022). As expected, salt treatment induced the expression of all three markers in Col-0 (wt) seedlings. Differences in marker gene induction were observed in some *sbt* lines compared with Col-0, although the overall effects were relatively modest (Figure 2A). We next assessed whether altered signaling was associated with changes in growth under saline conditions. Five-day-old seedlings were transferred to either mock or 100 mM NaCl-containing plates, and root responses were evaluated after 5 days by measuring primary root length and lateral root number (Figure 2B/C). No major differences in primary root length were observed between genotypes under salt conditions (Figure 2B/C). The *sbt5.3* mutant displayed shorter roots and fewer lateral roots already under control conditions. This phenotype is consistent with previous reports showing that *AIR3/SBT5.3* is specifically expressed at sites of lateral root emergence and contributes to lateral root development (Neuteboom et al., 1999). Under salt stress, most genotypes exhibited comparable primary root length, with no significant differences relative to salt treated wt seedlings (Figure 2B, S4A). Differences in lateral root traits were observed among the *sbt* mutants under salt treatment. In particular, *sbt1.1-3* produced fewer lateral roots than Col-0, whereas a significant reduction in lateral root density was detected only in *sbt1.6-2* (Figure 2C; Figure S4B). Analysis of cotyledon area revealed no major differences between genotypes under salt stress (Figure S4C), although some mutants, including *sbt1.1-1* and *sbt5*.3, displayed reduced fresh weight, already under control conditions (Figure S4D). Since ectopic lignin deposition is a hallmark of CW alteration and damage responses (Caño-Delgado et al., 2003; Denness et al., 2011; Engelsdorf et al., 2018; Van der Does et al., 2017), we next investigated whether loss of *SBT* affects salt-induced lignification. Six-day-old seedlings were treated with 200 mM NaCl for 48 h and stained with phloroglucinol to visualize lignin accumulation (Figure 2D). While mock-treated seedlings did not display detectable lignification, salt treatment induced ectopic lignin deposition in the root tip region. Quantification of lignified root areas revealed a significant increase in both *sbt1.1* alleles and in *sbt5.3* compared with salt treated wt seedlings (Figure 2D) suggesting that SBT1.1 and SBT5.3 may contribute to the regulation of CW remodeling during salt stress.

**Figure 2.**
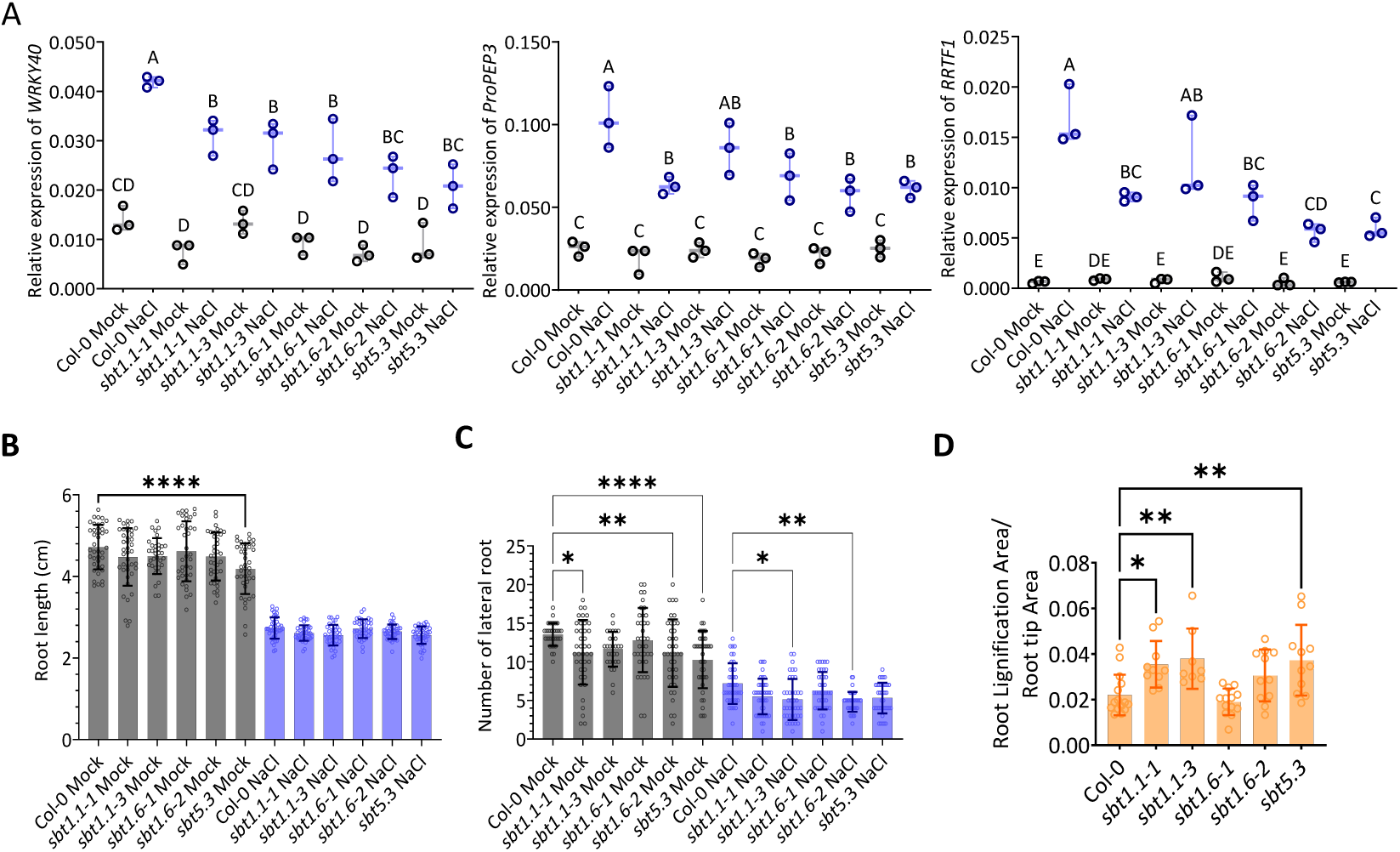
Early and late responses to salt stress in *sbt* mutant lines. (A) Expression analysis of salt-responsive marker genes. Six-day-old seedlings were treated with either mock medium or medium supplemented with 100 mM NaCl for 1 h. Expression of *WRKY40*, *ProPEP3*, and *RRTF1* was analyzed relative to *ACT2*, used as a reference gene. Three independent biological replicates were analyzed for each genotype and treatment, each consisting of 25 seedlings (n = 3). Data are presented as box-and-whisker plots showing the median and interquartile range (25^th^-75^th^ percentiles), with whiskers extending from the minimum to the maximum value. Each dot represents one independent biological replicate. Statistical significance was assessed using two-way ANOVA with genotype and treatment as factors, followed by Tukey’s HSD test for multiple comparisons. Groups labeled with different letters are significantly different (P < 0.05). (B) Primary root length and (C) lateral root number were measured in 10-day-old seedlings. Seedlings were grown for 5 days on mock medium and then transferred to either mock or 100 mM NaCl plates for an additional 5 days. Data were obtained from two independent experiments, each including 20 seedlings per genotype and treatment (n = 40 seedlings in total per genotype and treatment). Each dot represents an individual seedling. Data are presented as mean ± SD. Statistical significance was determined by comparing each mutant line with Col-0 within each treatment using one-way ANOVA with Šídák correction for multiple comparisons. Selected pairwise comparisons are shown. (D) Six-day-old seedlings were treated with either mock or 200 mM NaCl for 48 h and subsequently stained with phloroglucinol to visualize lignin deposition. The lignified area was quantified using Fiji-ImageJ and expressed relative to the total root tip area. The experiment was independently repeated three times with similar results. Data from one representative experiment are shown (n = 10 seedlings per genotype and treatment), with each dot representing an individual seedling. Data are presented as mean ± SD. Statistical significance was determined by comparing each mutant line with Col-0 under NaCl treatment using one-way ANOVA with Šídák correction for multiple comparisons. Unless otherwise indicated, significance levels are denoted as ns (P ≥ 0.05), * (P < 0.05), ** (P < 0.01), *** (P < 0.001), and **** (P < 0.0001).

### Application of the PSK peptide alleviates both early and late salt-triggered responses

Since SBT1.1 has been shown to be required for PSK4 maturation (Srivastava et al., 2008), and a direct connection between PSK signaling and salt stress responses has not yet been established, we further investigated the role of PSK signaling under salt stress. PSK precursors encode a highly conserved pentapeptide sequence, YIYTQ (Kaufmann and Sauter, 2019), which is sulfated by tyrosylprotein sulfotransferases (TPSTs). PSK is perceived by the Leucine-Rich Repeat Receptor-like Kinases PSKR1 and PSKR2 (Kaufmann et al., 2021). Given that PSK is widely present in spermatophytes and has previously been shown to promote root growth in Arabidopsis (Kutschmar et al., 2009; Li et al., 2024), we used exogenous PSK application together with the loss-of-function PSK receptor double mutant *pskr1-3 pskr2-1* (Kaufmann and Sauter, 2019; Sauter, 2015; Stührwohldt et al., 2021) in our salt stress experiments. To assess the effects of PSK signaling on both early and late salt stress responses, we performed a series of assays covering distinct stages of the response, including MPK6 phosphorylation (15 min), gene expression (1 h), lignification (48 h), and root growth (5 d) (Figure 3A). We first analyzed MPK6 phosphorylation as an early response to salt stress. Six-day-old seedlings were treated with mock, NaCl, PSK, or a combination of NaCl and PSK for 15 min. Salt-induced MPK6 phosphorylation was reduced in PSK-treated wt seedlings, whereas no effect was observed in the *pskr1-3 pskr2-1* mutant (Figure 3B, S5A/B). We next examined the expression of salt-induced marker genes (*WRKY40*, *ProPEP3*, and *RRTF1*). Under NaCl treatment, PSK application significantly reduced the expression of these marker genes in wt, while no significant effect was observed in the *pskr1-3 pskr2-1* mutant (Figure 3C), indicating that PSK attenuates salt-induced signaling in a PSKR-dependent manner. Notably, in the absence of PSK treatment, *pskr1-3 pskr2-1* mutant showed reduced induction of *WRKY40* and *RRTF1* upon salt stress, whereas *ProPEP3* induction remained comparable to that observed in wt seedlings. These results may indicate that PSKR differentially contributes to salt-induced transcriptional responses, while PSK-mediated attenuation of these responses requires functional PSKR receptors. To assess the impact of PSK on salt-induced phenotypes, we first quantified salt-induced ectopic lignin accumulation. Six-day-old seedlings were treated with NaCl for 48 h in the presence/absence of PSK. In wt seedlings, PSK application significantly reduced salt-induced ectopic lignin deposition (Figure 3D). In contrast, the *pskr1-3 pskr2-1* mutant displayed higher salt-induced lignin accumulation compared to wt and this phenotype was not alleviated by PSK treatment (Figure 3D). Next, seedlings were treated with mock and PSK in the presence or absence of NaCl for 5 days (Figure 3E). Although PSK has previously been reported to promote primary root growth in Arabidopsis (Kutschmar et al., 2009), under our experimental conditions PSK treatment did not significantly increase root length compared with the mock control. Under salt stress, however, PSK application alleviated NaCl-induced root growth inhibition in wt (Figure 3E, S5B), whereas *pskr1-3 pskr2-1* exhibited no response to PSK (Figure 3E), indicating that PSKR-dependent PSK signaling contributes to the attenuation of salt-induced lignification and growth inhibition.

**Figure 3.**
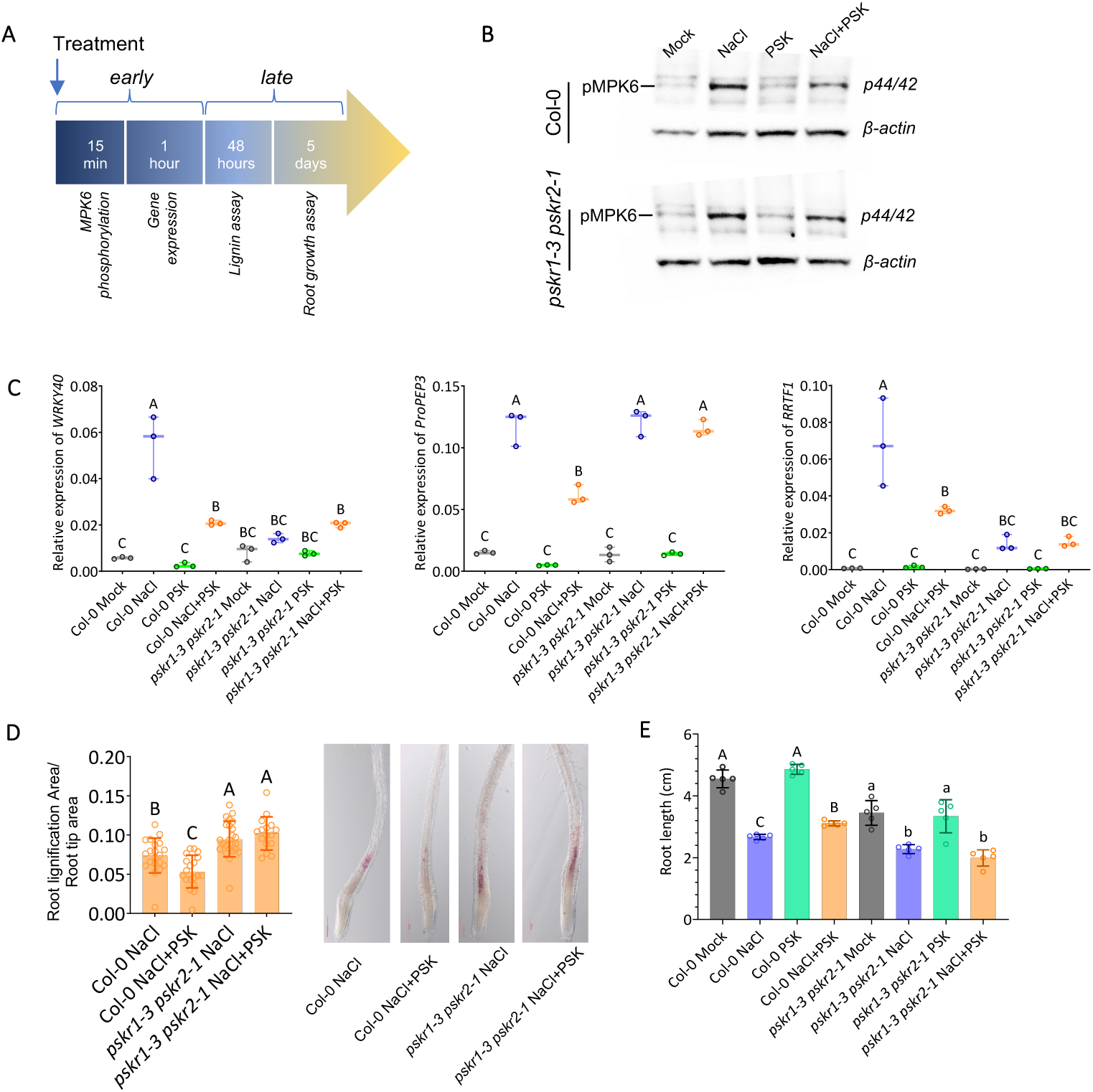
Application of PSK-α attenuates salt-triggered signaling and physiological responses. (A) Timeline of the experiments used to assess early and late responses to salt stress and PSK-α treatment. (B) MPK phosphorylation analysis in Col-0 and *pskr1-3 pskr2-1*. Six-day-old seedlings were treated for 15 min with mock medium or medium supplemented with 100 mM NaCl, 1 μM PSK-α, or a combination of 100 mM NaCl and 1 μM PSK-α. Phosphorylation was detected by immunoblotting using an anti-phospho-*p44/42* MAPK antibody (upper panel), with an anti β-actin antibody used as a loading control (lower panel). The experiment was independently repeated twice with similar results. (C) Expression analysis of salt-responsive marker genes. Six-day-old seedlings were treated for 1 h with mock medium or medium supplemented with 100 mM NaCl, 1 μM PSK-α, or a combination of 100 mM NaCl and 1 μM PSK-α. Three independent biological replicates were analyzed for each genotype and treatment, each consisting of 25 seedlings (n = 3). Each dot represents one independent biological replicate. Data are presented as box-and-whisker plots showing the median and interquartile range (25^th^-75^th^ percentiles), with whiskers extending to the minimum and maximum values. Data were analyzed by two-way ANOVA with genotype and treatment as factors, followed by Tukey’s HSD post hoc test. Groups labeled with different letters are significantly different (P < 0.05). (D) Six-day-old seedlings were treated for 48 h with mock medium, medium supplemented with 200 mM NaCl, or medium supplemented with 200 mM NaCl and 1 μM PSK-α. Roots were stained with phloroglucinol to visualize lignin deposition. Lignified areas were quantified using Fiji-ImageJ and expressed relative to the total root tip area. Each dot represents an individual seedling (n = 20) from two independent experiments. Data are presented as mean ± SD. Statistical analysis was performed using one-way ANOVA followed by Tukey’s HSD test (P < 0.05). Representative images are shown. (E) Five-day-old Col-0 and *pskr1-3 pskr2-1* seedlings were transferred to control plates or plates supplemented with 100 mM NaCl, 1 μM PSK-α, or a combination of 100 mM NaCl and 1 μM PSK-α. Primary root length was measured after 5 d of treatment. The experiment was independently repeated three times with similar results. Data from one representative experiment are shown (n = 5 seedlings per genotype and treatment), with each dot representing an individual seedling. Data are presented as mean ± SD. Statistical significance was determined using one-way ANOVA followed by Tukey’s HSD test to compare treatments within each genotype. Uppercase letters indicate significant differences among treatments in Col-0, whereas lowercase letters indicate significant differences among treatments in *pskr1-3 pskr2-1* (P < 0.05).

### PSK limits salt-induced changes in cell wall monosaccharide composition

Given that salt stress induces CW remodeling, and that PSK has been associated with changes in CW composition during protoplast regeneration (Godel-Jędrychowska et al., 2019), we analyzed the monosaccharide composition of alcohol-insoluble residue (AIR) extracted from 6-day-old seedlings treated for 48 h with NaCl, PSK, or a combination of both. As previously reported by (Yan et al., 2021; Zhang et al., 2023) NaCl treatment increased the AIR-normalized content of several monosaccharides, including fucose, rhamnose, arabinose, galactose, mannose, and xylose (Figure 4). A similar trend was observed in the *pskr1-3 pskr2-1* mutant, although mannose and xylose levels did not differ significantly between NaCl-treated and control seedlings. Upon PSK co-treatment, wt seedlings displayed attenuated salt-induced increases in the AIR-normalized content of several monosaccharides, particularly rhamnose, arabinose, and mannose, whereas galactose levels remained largely unchanged (Figure 4). In contrast, these effects were not observed in the *pskr1-3 pskr2-1* mutant, suggesting that PSK-dependent modulation of AIR monosaccharide profiles requires functional PSK signaling. Analysis of monosaccharide molar percentages revealed a more selective effect on CW composition. Among the monosaccharides analyzed, arabinose showed the most pronounced salt-induced change in relative abundance, which was attenuated by PSK in Col-0 but not in the *pskr1-3 pskr2-1* mutant (Figure S6).

**Figure 4.**
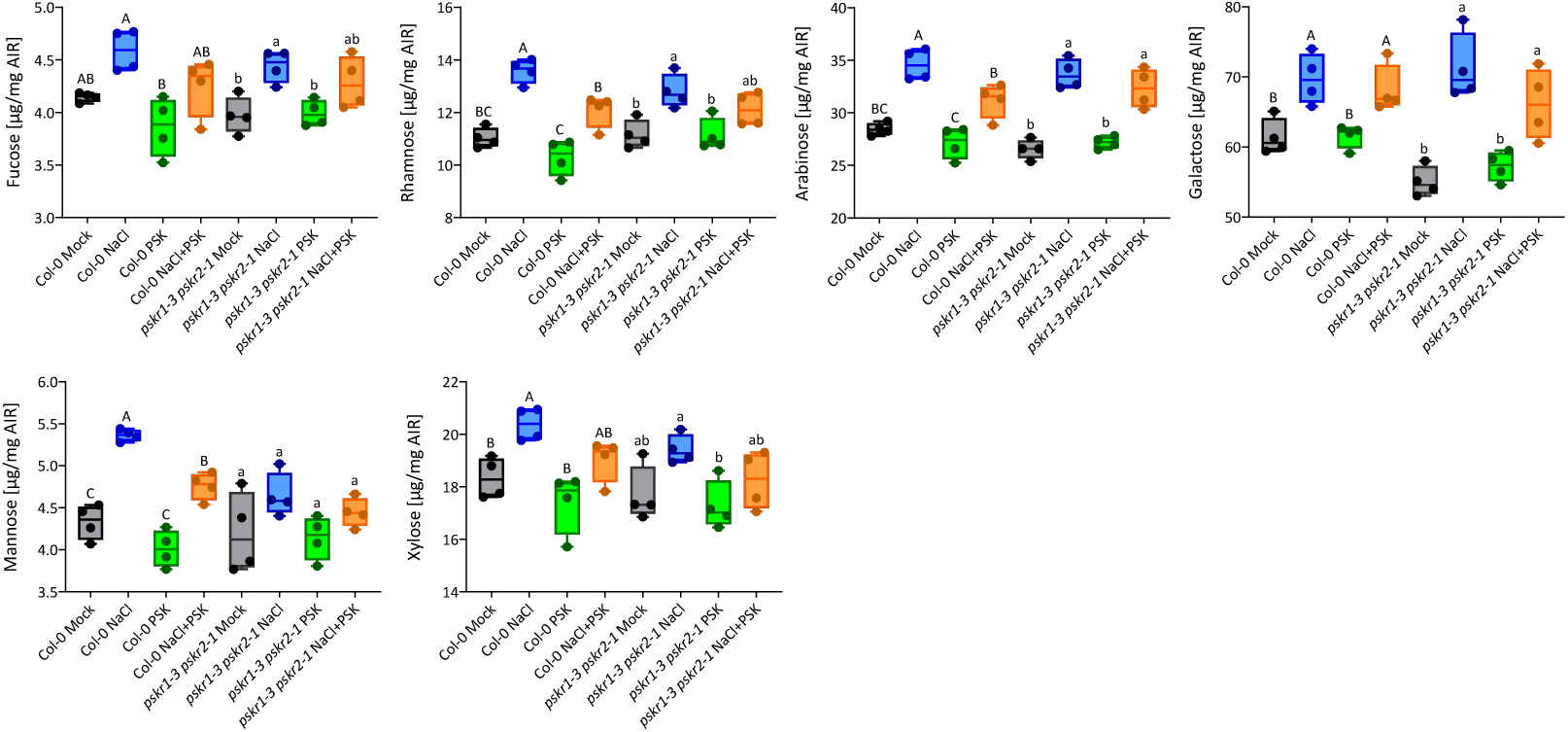
PSK-α attenuates NaCl-induced changes in cell wall composition. Six-day-old Col-0 and *pskr1-3 pskr2-1* seedlings were treated for 48 h with mock medium, 100 mM NaCl, 1 μM PSK-α, or a combination of 100 mM NaCl and 1 μM PSK-α. Quantification of cell wall-associated monosaccharides in Col-0 and *pskr1-3 pskr2-1* under NaCl treatment in the presence or absence of PSK-α. Fucose, rhamnose, arabinose, galactose, mannose, and xylose were quantified and expressed as µg mg⁻¹ alcohol-insoluble residue (AIR). Four independent biological replicates were analyzed for each genotype and treatment, each consisting of 40 seedlings (n = 4). Each dot represents one independent biological replicate. Data are presented as box-and-whisker plots showing the median and interquartile range (25^th^-75^th^ percentiles), with whiskers extending from the minimum to the maximum value. Statistical analysis was performed using one-way ANOVA followed by Tukey’s HSD test to compare treatments within each genotype (P < 0.05). Different uppercase letters indicate significant differences among treatments in Col-0, while different lowercase letters indicate significant differences among treatments in *pskr1-3 pskr2-1*.

### PSK attenuates salt stress signaling independently of cell wall integrity receptors but requires FER signaling for adaptive root growth

As PSK attenuates NaCl-induced changes in CW composition, we next asked whether its effects on salt stress responses depend on CWI perception. To address this question, we analyzed the response to PSK in the CWI mutants *fer-4* and *herk1 the1-4*, which display enhanced sensitivity to salt stress (Feng et al., 2018; Gigli-Bisceglia et al., 2022). To examine early signaling events, seedlings were treated for 15 min with mock, NaCl, PSK, or NaCl in combination with PSK, and MPK phosphorylation was analyzed (Figure 5A). NaCl induced MPK6 phosphorylation in Col-0, with a stronger activation in both CWI mutants (Gigli-Bisceglia et al., 2022). PSK co-treatment reduced NaCl-induced MPK6 phosphorylation in all genotypes, indicating that attenuation of early salt signaling by PSK occurs independently of FER-, HERK1- and THE1-mediated CWI signaling. To further examine downstream responses, we analyzed the expression of salt-responsive marker genes. Consistent with the MPK analysis, PSK reduced NaCl-induced *ProPEP3* expression in all genotypes except *pskr1-3 pskr2-1* (Figure 5B), demonstrating that repression of early salt signaling depends on PSKR but not on the CWI receptors analyzed. We next investigated whether attenuation of early signaling translated into improved growth under salt stress. In Col-0, PSK partially restored primary root growth in the presence of NaCl, whereas no significant growth recovery was observed in either *fer-4* or *herk1 the1-4* (Figure 5C). Instead, PSK induced marked changes in root morphology under saline conditions, most prominently in *fer-4*, where NaCl and PSK together triggered a pronounced root bending phenotype (Figure 5D). Thus, although PSK is able to dampen early salt stress signaling independently of FER-, HERK1- and THE1-mediated CWI perception, proper CWI signaling appears to be required to translate these signaling outputs into coordinated root growth. In the absence of FER-dependent CWI sensing, PSK-mediated growth responses become aberrant, resulting in altered root growth orientation.

**Figure 5.**
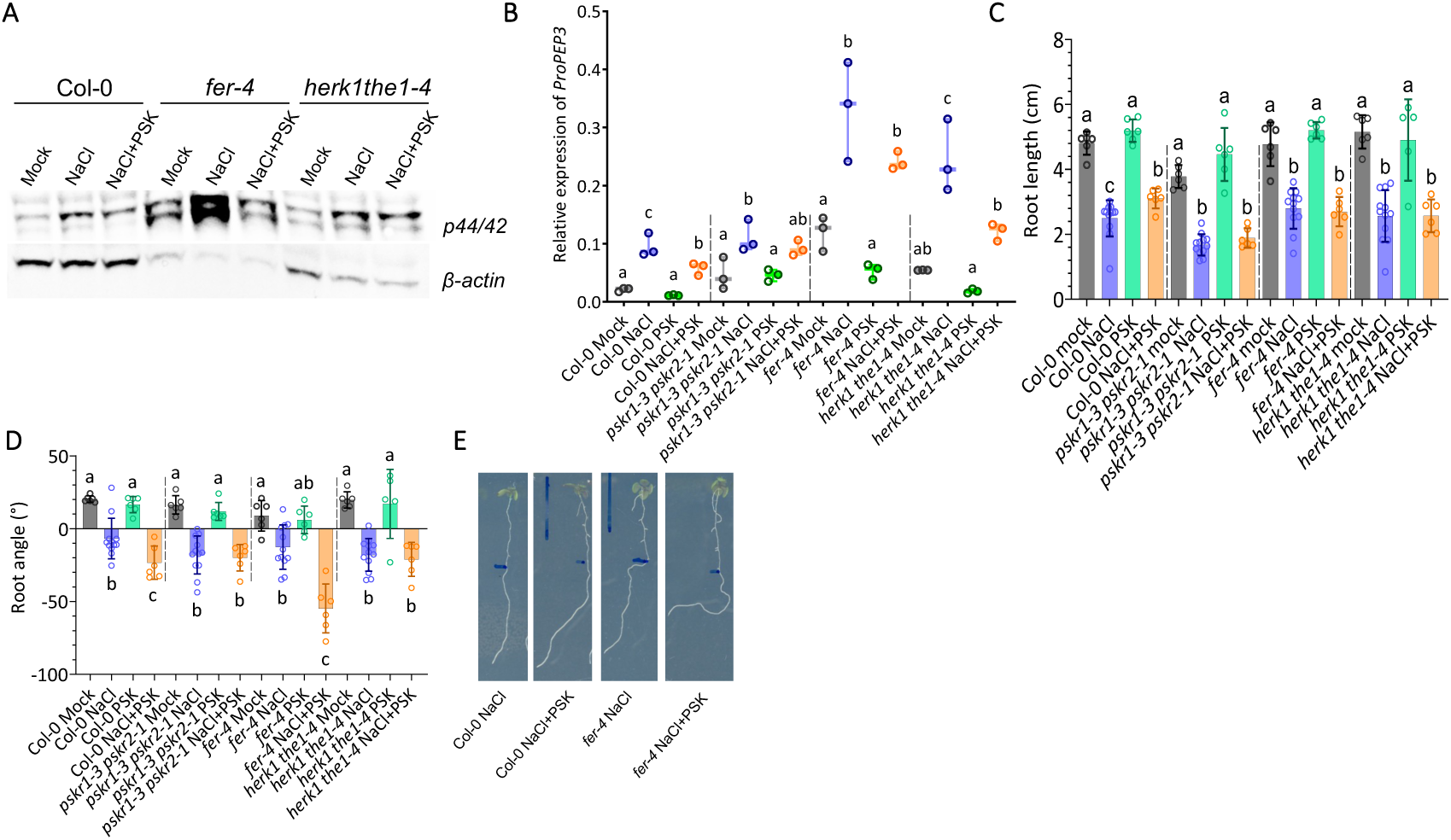
PSK-α attenuates early salt stress signaling but does not alleviate salt-induced root growth inhibition in CWI receptor mutants. (A) MPK phosphorylation analysis in six-day-old Col-0, *fer-4*, and *herk1 the1-4* seedlings treated for 15 min with mock medium or medium supplemented with 100 mM NaCl, 1 μM PSK-α, or a combination of 100 mM NaCl and 1 μM PSK-α. β-actin was used as a loading control. The experiment was independently repeated twice with similar results, and one representative immunoblot is shown. (B) *ProPEP3* expression analysis in 6-day-old seedlings treated for 1 h with mock medium or medium supplemented with 100 mM NaCl, 1 μM PSK-α, or a combination of NaCl and PSK-α. Three independent biological replicates were analyzed for each genotype and treatment, each consisting of 25 seedlings (n = 3). Each dot represents one independent biological replicate. Data are presented as box-and-whisker plots showing the median and interquartile range (25^th^-75^th^ percentiles), with whiskers extending from the minimum to the maximum value. (C) Primary root growth and (D) root skewing in Col-0, *fer-4*, and *herk1 the1-4*. Five-day-old seedlings were transferred to MS/2 medium supplemented with 0.5% (w/v) sucrose and either 100 mM NaCl, 1 μM PSK-α, or a combination of 100 mM NaCl and 1 μM PSK-α, with non-supplemented medium used as the mock control. Sample sizes were n = 6 seedlings per genotype for the mock, PSK-α, and NaCl + PSK-α treatments and n = 12 seedlings per genotype for the NaCl treatment. Each dot represents an individual seedling. Data are presented as mean ± SD. (E) Representative images of Col-0 and *fer-4* seedlings grown under NaCl or NaCl + PSK-α conditions, illustrating the root growth and skewing phenotypes quantified in (C) and (D). Statistical significance was determined using one-way ANOVA followed by Tukey’s HSD test to compare treatments within each genotype (P < 0.05)

## Discussion

Salt stress induces extensive remodeling of the plant cell wall, including changes in polysaccharide composition, pectin methylesterification, and ectopic lignin deposition (Colin et al., 2023; Gigli-Bisceglia et al., 2022; Sharma et al., 2026; Zhang et al., 2023; Zou et al., 2022). While several plasma membrane-localized receptors and intracellular signaling components involved in these processes have been described (Debnath et al., 2026), the contribution of apoplast-localized proteins to salt stress responses remains less well understood. In this study, we addressed this gap by performing an apoplastic proteomic analysis and identified a set of proteins whose abundance is altered upon NaCl treatment. Among them, we identified several proteins belonging to the subtilisin-like protease (SBT) family, including SBT1.1, SBT1.6, and SBT5.3. Although the corresponding mutant lines exhibited relatively mild phenotypes under salt stress, with some showing significantly different growth parameters under control conditions, both *sbt1.1* and *sbt5.3* showed increased accumulation of ectopic lignin upon salt treatment. SBT1.1 is of particular interest, as it has been previously shown to process the precursor of the sulfated peptide phytosulfokine (PSK4) (Srivastava et al., 2008). This observation prompted us to investigate whether PSK signaling might play a role in modulating salt stress responses. PSK has been associated with growth regulation through cell expansion and with drought stress responses (Kutschmar et al., 2009; Ladwig et al., 2015; Stührwohldt et al., 2021), however its function in early signaling and CW responses to salinity remains poorly understood. While our experiments employed the mature sulfated PSK pentapeptide, the distinct transcriptional responses of individual PSK precursor genes to salt stress suggest that endogenous PSK signaling is likely regulated in a gene-specific manner. Thus, the effects observed following exogenous PSK application may represent only one component of the endogenous PSK signaling network. In our study, exogenous application of PSK attenuated early salt-induced signaling responses, as reflected by reduced MPK6 phosphorylation and decreased expression of salt-responsive marker genes in wild-type seedlings. Importantly, MPK6 phosphorylation (Gigli-Bisceglia et al., 2022; Ichimura et al., 2000), *RRTF1* induction (Lamers et al., 2025) and ectopic lignin deposition (Sharma et al., 2026) are characteristic features of salt stress and are not typically induced to the same extent by purely osmotic treatments. These effects were not observed in the *pskr1-3 pskr2-1* mutant, indicating that PSK-mediated modulation of salt-induced signaling requires functional PSK receptors. Interestingly, in the absence of exogenous PSK, the *pskr1-3 pskr2-1* mutant already displayed reduced induction of a subset of marker genes upon salt treatment, while exhibiting increased ectopic lignin accumulation under salt stress, a characteristic feature of CW damage responses associated with CW perturbation that is not generally observed under osmotic treatment (Engelsdorf et al., 2018). Together, these observations suggest that PSK signaling contributes to shaping a specific branch of the salt stress response. The fact that these salt-associated outputs are modulated by PSK further suggests that PSK signaling may specifically interfere with the ionic- or damage-associated component of the salt response, rather than acting only through a general osmotic stress pathway. This interpretation is consistent with the observed reduction in ectopic lignin deposition and the partial restoration of root growth in PSK-treated Col-0 plants under salt stress, both of which were absent in the PSK receptor mutants.

Salt treatment led to an increased accumulation of several CW-associated monosaccharides, consistent with previous reports describing extensive remodeling of the primary CW under salinity (Colin et al., 2023; Gigli-Bisceglia et al., 2022; Zhang et al., 2023; Zou et al., 2024). These changes likely reflect modifications of wall polymers involved in stress adaptation. The increased abundance of rhamnose and arabinose is consistent with remodeling of the rhamnogalacturonan-I (RG-I) domain of pectins, whose arabinan-rich side chains have been implicated in maintaining wall flexibility and integrity during abiotic stress (Debnath et al., 2026). Arabinose may also derive from other wall-associated glycoconjugates, including arabinogalactan proteins (AGPs) (Lamport et al., 2006; Tan et al., 2024) and hydroxyproline-rich glycoproteins (Zou et al., 2024), both of which have been linked to salt-induced CW remodeling. PSK co-treatment attenuated the salt-induced accumulation of rhamnose, arabinose and mannose observed in Col-0, whereas these effects were absent in the *pskr1-3 pskr2-1* mutant. This indicates that modulation of CW composition by PSK requires functional PSKR signaling. Transcriptomic analyses of PSK-responsive seedlings revealed an enrichment of genes associated with CW modification, CW organization and pectin metabolism (Kaufmann et al., 2021), suggesting a broader role for PSK signaling in regulating CW dynamics. To further understand how PSK-mediated modulation of salt stress responses is integrated with CWI signaling, we investigated its effects in mutants impaired in CWI perception. In both *fer-4* and *herk1 the1-4*, PSK attenuated NaCl-induced MPK6 phosphorylation, indicating that repression of early salt signaling occurs independently of the analyzed CWI receptors. Although PSK did not significantly restore root growth under salt stress in either mutant, the enhanced salt sensitivity of these genotypes (Feng et al., 2018; Gigli-Bisceglia et al., 2022) makes it difficult to conclude whether this reflects a specific requirement for CWI signaling in PSK-mediated growth promotion. Instead, the most striking phenotype was observed in *fer-4*, where PSK treatment under salt stress induced a pronounced root bending response. This observation suggests that, although PSK signaling remains active in the absence of FER-dependent CWI perception, its developmental effect on roots is no longer properly coordinated in the absence of this receptor. Given that FER has recently been shown to regulate pectin homeostasis through the control of pectin methylesterase activity (Biermann et al., 2025), the altered pectin environment in *fer-4* may influence how PSK-mediated changes in CW remodeling are translated into adaptive growth responses. Thus, CWI signaling does not appear to be required for PSK-mediated attenuation of early salt responses, but rather for directing these signaling outputs into an appropriate developmental response. Together, our findings suggest that PSK signaling acts as a modulator of salt-induced responses, influencing both early signaling outputs and downstream changes in CW composition.

## Materials and Methods

### Plant materials and growth conditions

*Arabidopsis thaliana* ecotype Columbia-0 (Col-0) was used as the wild type in all experiments. The mutant lines used in this study are listed in Table S3. Genotypes were confirmed by PCR using genomic DNA extracted from adult leaves with the Phire Tissue Direct PCR Kit (Thermo Scientific). Previously published primer sets were used when available, whereas primers designed in this study are listed in Table S3. Seeds were surface-sterilized in 20% (v/v) commercial bleach for 10 min, rinsed four times with sterile Milli-Q water, and stratified in the dark at 4°C for 2 d before sowing. For time-course and *SBT* gene-expression analyses, seedlings were grown for 6 d in 250-mL Erlenmeyer flasks containing 110 ml half-strength Murashige and Skoog liquid medium (MS/2), supplemented with vitamins and MES (2.5 g L⁻¹), and 1% (w/v) sucrose, adjusted to pH 5.7. The growth medium was then replaced with the corresponding treatment medium. For MPK phosphorylation, gene-expression, and cell-wall composition analyses, seedlings were grown in 12-well plates containing 2 mL per well of liquid MS/2 medium supplemented with vitamins and MES (2.5 g L⁻¹), and 0.5% (w/v) sucrose, adjusted to pH 5.7. After 5 d of growth, the medium was replaced with fresh medium, and treatments were applied on day 6. For lignin analyses, seedlings were grown in liquid MS/2 medium supplemented with vitamins and MES (2.5 g L⁻¹) and 1% (w/v) sucrose, adjusted to pH 5.7. Unless otherwise indicated, seedlings used for MPK phosphorylation, gene-expression, lignin-staining, and cell-wall analyses were treated at 6 d after germination. Root-growth assays were performed on vertically oriented square plates containing MS/2 medium supplemented with vitamins and MES (2.5 g L⁻¹), 0.5% (w/v) sucrose, and 1% (w/v) Daishin agar (Duchefa). Seedlings were grown vertically for 5 d, transferred to treatment plates, and grown for additional 5 days before root length was measured. Unless otherwise indicated, treatments consisted of Milli-Q water as the mock control, 100 mM NaCl, 1 μM synthetic phytosulfokine (PSK; Y(SO₃H)-I-Y(SO₃H)-T-Q; Peptanova, catalog no. 4477-s), or a combination of NaCl and PSK. For lignin staining, seedlings were treated with 200 mM NaCl. All plants were grown under long-day conditions consisting of 16 h light at 22°C and 8 h darkness at 18°C, with a light intensity of 130 μmol m⁻² s⁻¹.

### Apoplastic protein extraction and proteomic analysis

Three biological replicates, each consisting of 30 mg of Arabidopsis thaliana Col-0 seeds, were grown in 250-mL Erlenmeyer flasks containing 110 mL of MS/2 liquid medium supplemented with vitamins, MES (2.5 g L⁻¹), 1% (w/v) sucrose, and adjusted to pH 5.7. Seedlings were grown for 6 d under long-day conditions (16 h light/8 h dark) at 21°C with continuous shaking (130 rpm). The growth medium was then replaced with fresh medium in the presence or absence of 100 mM NaCl, and seedlings were incubated for an additional 24 h. Following treatment, seedlings and culture media were processed separately. Seedlings were gently washed three times with sterile Milli-Q water and transferred to 6-well plates containing 6 mL of infiltration buffer (IB), (50 mM KH₂PO₄, 0.2 M KCl, 1 mM PMSF, and 1× protease inhibitor cocktail (Sigma-Aldrich, P9599), pH 6.2). Seedlings were vacuum infiltrated with the IB for 10 min before being transferred to syringes without plungers. Apoplastic extracts were recovered by centrifugation at 600 × g for 25 min. Growth media were supplemented with 1× protease inhibitor cocktail prior to further processing. Apoplastic extracts and growth media were subsequently enriched by using the FASP method with 10-kDa molecular weight cut-off centrifugal filters. After overnight trypsin digestion, tryptic peptides were eluted with 50 mM ammonium bicarbonate, acidified and desalted using in-house made C18 spin tips. Samples were measured on an Exploris 480 (Thermo Scientific) using data dependent acquisition as previously described (Kuhn et al., 2024). The MaxQuant quantitative proteomics software package was used to analyse LC–MS data with all MS/MS spectra (Tyanova et al., 2016a). The following settings were used: peptide and protein FDR ≤ 0.01; the proteome of *A. thaliana* (UniProt ID UP000006548) was used as the protein database; maximum missed cleavage was set at 2; variable modifications Oxidation (M), Acetyl (protein N-term), Deamidation (NQ), fixed modification AcrylAmide (C); match between runs and label-free quantification options were selected (Kuhn et al., 2024). The proteinGroups file was analyzed using the Perseus software (Tyanova et al., 2016b). Data were filtered for reverse, contaminants and only identified by site. MaxLFQ intensity values were log_2_ transformed and filtered to contain at least 75% valid values in each group. Values were subsequently normalized by median column subtraction. Remaining missing values were imputed from a normal distribution using standard settings in Perseus (width: 0.3, down shift: 1.8). An FDR permutation-based t-test was performed to identify significantly changing proteins (FDR ≤0.05).

### MPK phosphorylation assay and gene expression analysis

Six-day-old seedlings were grown as described above and treated with mock, 100 mM NaCl, 1 μM PSK, or a combination of NaCl and PSK for 15 min (MPK phosphorylation assay) or 1 h (gene expression analysis). Following treatment, seedlings were flash-frozen in liquid nitrogen, transferred to 2-mL tubes containing a sterile stainless-steel bead, and homogenized using a TissueLyser (Qiagen). For gene expression analysis, total RNA was extracted using the EasyPrep Plant Total RNA Extraction Miniprep Kit (Bioland) according to the manufacturer’s instructions. Quantitative RT-PCR was performed using a CFX96 Real-Time PCR Detection System (Bio-Rad) with qPCRBIO SyGreen Blue Mix Lo-ROX (Sopachem) and transcript-specific primers (Table S3). Relative transcript levels were calculated using *ACTIN2* as the reference gene, as previously described (Engelsdorf et al., 2018; Gigli-Bisceglia et al., 2022, 2018). For MPK phosphorylation analysis, proteins were extracted in ice-cold extraction buffer (50 mM Tris-HCl, pH 7.5, 200 mM NaCl, 1 mM EDTA, 10% glycerol, 0.1% Tween-20, 1 mM PMSF, 1 mM DTT, 1× protease inhibitor cocktail and 1× phosphatase inhibitor cocktail (P2850, Sigma-Aldrich)). Samples were incubated on ice for 15 min and centrifuged at 13,000 × *g* for 15 min at 4°C. Protein concentration was determined using the Bradford assay (Bio-Rad), and 25 μg of total protein was separated by 10% precast SDS-PAGE gels (Bio-Rad) and transferred onto nitrocellulose membranes using the Trans-Blot Turbo Transfer System (Bio-Rad). Membranes were blocked for 2 h in TBST containing 5% (w/v) bovine serum albumin and incubated overnight at 4°C with anti-phospho-p44/42 MAPK (Erk1/2) antibody (Thr202/Tyr204; Cell Signaling Technology, #9101; 1:2000). After washing, membranes were incubated for 2 h with HRP-conjugated anti-rabbit IgG (Cell Signaling Technology, #7074; 1:6000). Signals were detected using Clarity Western ECL Substrate (Bio-Rad) and visualized with a ChemiDoc Imaging System (Bio-Rad). Equal loading was verified after membrane stripping by probing with anti-β-actin-HRP linked antibody (Santa Cruz Biotechnology, sc-47778; 1:2000).

### Lignin staining and quantification

Six-day-old seedlings were treated with 200 mM NaCl in the presence or absence of 1 μM PSK for 48 h, fixed in 70% (v/v) ethanol for 3 days, and stained with freshly prepared phloroglucinol solution (0.1% (w/v) phloroglucinol in 20% (v/v) HCl) to visualize lignin (Engelsdorf et al., 2018; Gigli-Bisceglia et al., 2018). Images of the root tip were acquired using a Zeiss Axio Zoom.V16 fluorescence stereomicroscope at 63× magnification. Lignin accumulation was quantified in Fiji/ImageJ by measuring the lignified and total root tip areas, and is presented as the ratio of lignified to total root tip area as in (Engelsdorf et al., 2018).

### Cell wall composition analysis

Six-day-old seedlings were treated for 48 h with MS/2 liquid medium supplemented with vitamins, MES (2.5 g L⁻¹), and 1% (w/v) sucrose, and adjusted to pH 5.7. NaCl (100 mM) and PSK (1 μM) were dissolved directly in the culture medium and applied either individually or in combination. Following treatment, seedlings were flash-frozen in liquid nitrogen, freeze-dried, and processed to obtain alcohol-insoluble residue (AIR) as previously described (Zhang et al., 2023; Zou et al., 2024). Briefly, freeze-dried tissue was sequentially washed with 70% (v/v) ethanol, chloroform:methanol (1:1, v/v), and acetone before drying overnight to obtain AIR. Starch was removed as described by (Pettolino et al., 2012). Cell wall monosaccharide composition was determined from 1-2 mg de-starched AIR hydrolyzed in 4% (w/v) sulfuric acid by autoclaving at 121°C for 60 min. Monosaccharides were diluted with ultrapure water, and ribose was added as an internal standard. Analysis was performed via high-performance anion-exchange chromatography with pulsed amperometric detection (HPAEC-PAD) using a biocompatible Knauer Azura HPLC system and an Antec Decade Elite SenCell detector heated to 40°C as described (Reckleben et al., 2025). Monosaccharides were separated using a Dionex CarboPac PA20 analytical column (Thermo Fisher Scientific), and individual monosaccharides were quantified using pure standards. The cell wall neutral monosaccharide composition represents the molar percentage of fucose, rhamnose, arabinose, galactose, xylose and mannose in relation to the sum of all neutral monosaccharides.

### Root growth assay and phenotyping

Five-day-old seedlings were transferred to MS/2 agar plates supplemented with 100 mM NaCl, 1 μM PSK, or a combination of NaCl and PSK, with non-supplemented medium used as the mock control. The position of the primary root tip was marked at the time of transfer, and plates were scanned after 5 days of treatment using an Epson Perfection V850 Pro flatbed scanner. Primary root elongation after transfer and lateral root number were quantified using RootNav software (Pound et al., 2013). Root skewing was quantified using SmartRoot 2.0 in Fiji. Root trajectories were traced from the root base to the root tip, and vector length was determined as the linear distance between these two points and used for root skewing analysis (Lobet et al., 2011). Lateral root density was calculated by dividing the number of lateral roots by the primary root length (Zhang et al., 2023). For shoot phenotyping, seedlings were harvested after 7 d of treatment. Cotyledon area was quantified from digital images using a custom image analysis pipeline, which is available upon request. Fresh weight was determined immediately after harvesting.

### Statistical Analysis

Statistical analyses and graph preparation were performed using GraphPad Prism v10.2.1 unless otherwise specified. Statistical significance is indicated by different letters or asterisks, as specified in the figure legends.

## Author contributions

Conceptualization: N.G.B.; Investigation: J.D., Z.J., T.v.d.M., N.B., M.R., C.T., T.E., and N.G.B.; Funding acquisition: T.E., C.T., and N.G.B.; Project administration: N.G.B.; Supervision: N.G.B.; Writing original draft: N.G.B.; Writing review and editing: All authors.

## Acknowledgments

We thank the late Prof. Dr. Sauter and Dr. Rehders (Plant Developmental Biology and Physiology, Christian-Albrechts-Universität Kiel, Germany) for kindly providing the *pskr1-3 pskr2-1* mutant seeds. This work was supported by the Dutch Research Council (NWO) through grants OCENW.M.24.079 and OCENW.XS23.1.050 awarded to N.G.B., by the Deutsche Forschungsgemeinschaft (DFG, German Research Foundation) grant EN 1071/3-1 awarded to T.E, and by the European Research Council (ERC) through the European Union’s Horizon 2020 Research and Innovation program (ERC Consolidator Grant agreement 724321) awarded to C.T.

