## Supplemental Figures for "Phytosulfokine signaling modulates salt stress responses and cell wall remodeling in Arabidopsis"

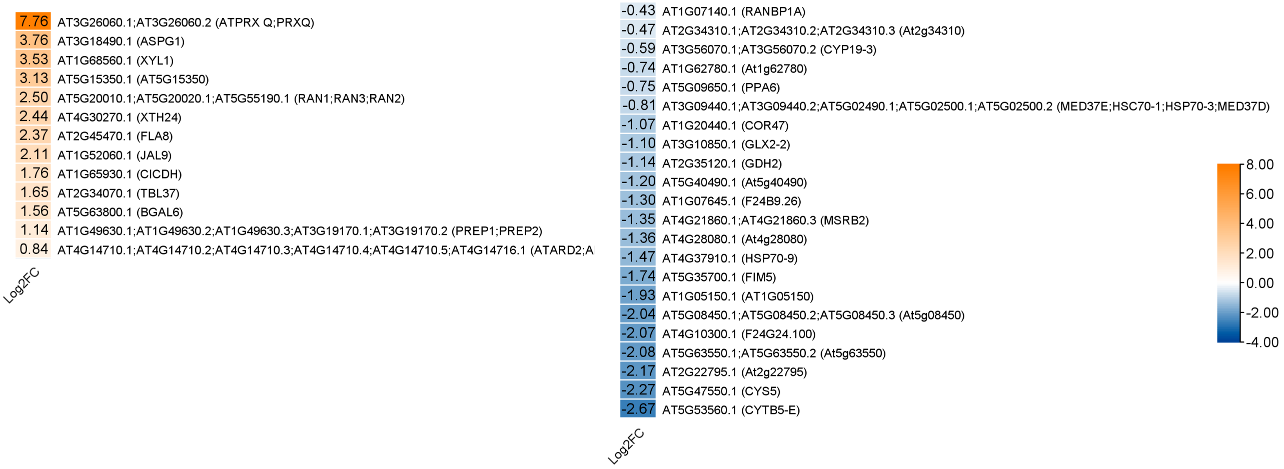


**Figure S1. Apoplastic proteomic analysis of Arabidopsis seedlings under salt stress.** Heatmap displaying the $log_2$ fold change of differentially accumulated proteins in the medium fraction of Col-0 seedlings treated with 100 mM NaCl for 24 h compared with mock-treated controls. Three independent biological replicates were analyzed per treatment, each consisting of approximately 1 g of seedlings (n = 3).

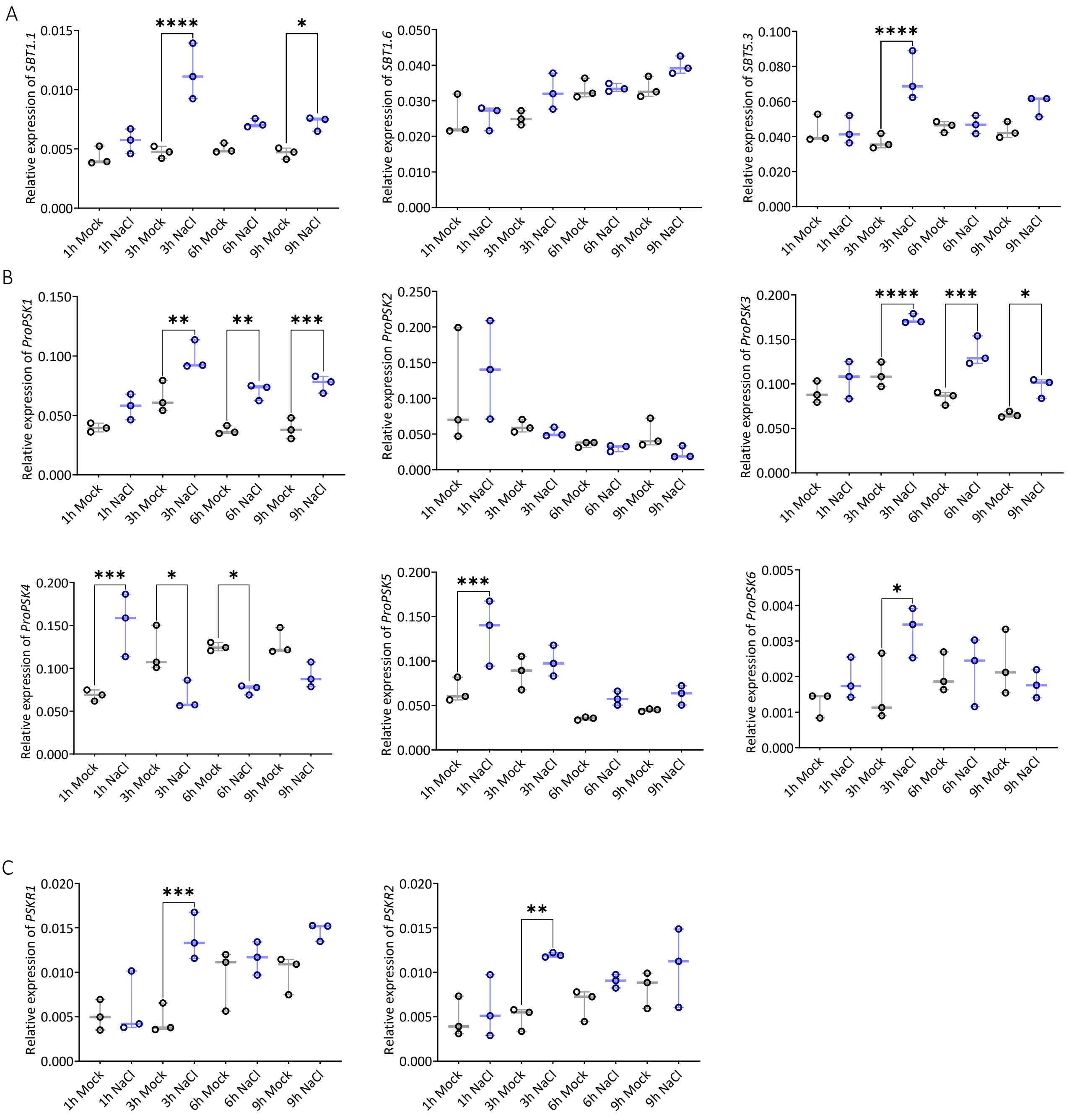


**Figure S2. Time-course expression analysis of candidate genes and PSK signaling components under salt stress.** (A) Time-course expression analysis of candidate genes identified by apoplastic proteomic analysis in six-day-old Col-0 seedlings treated with mock medium or medium supplemented with 100 mM NaCl. Samples were collected at 1, 3, 6, and 9 h after treatment. (B) Expression analysis of *ProPSK1*, *ProPSK2*, *ProPSK3*, *ProPSK4*, *ProPSK5*, and *ProPSK6* and (C) *PSKR1* and *PSKR2* in six-day-old Col-0 seedlings treated with mock medium or medium supplemented with 100 mM NaCl for the indicated times. Three independent biological replicates were analyzed for each treatment and time point, with each replicate consisting of 25 seedlings (n = 3). Each dot represents an independent biological replicate. Data are presented as box-and-whisker plots showing the median and interquartile range (25$^{th}$-75$^{th}$ percentiles), with whiskers extending from the minimum to the maximum value. Statistical significance was assessed using one-way ANOVA followed by Šídák's multiple-comparisons test to compare mock- and NaCl-treated seedlings at each time point. Significance levels are indicated as ns ($P \geq 0.05$), * ($P < 0.05$), ** ($P < 0.01$), *** ($P < 0.001$), and **** ($P < 0.0001$).

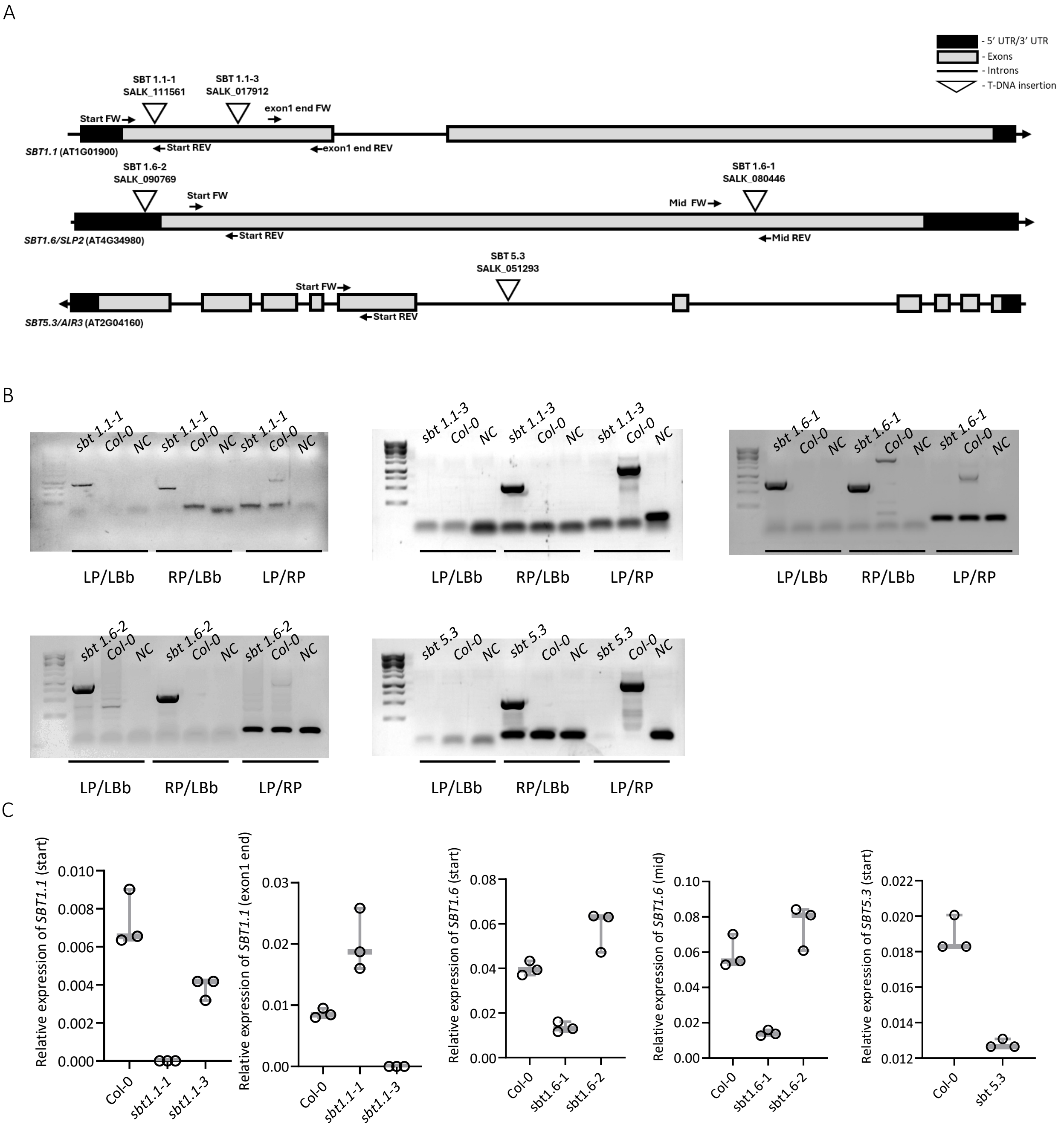


**Figure S3. Molecular characterization of T-DNA insertion lines.** (A) Schematic representation of T-DNA insertion sites and qRT-PCR primer positions for the candidate genes. Black boxes represent the 5′ and 3′ UTRs, grey boxes represent exons, black lines represent introns, and triangles indicate T-DNA insertions. (B) PCR-based genotyping of the candidate T-DNA insertion lines. Representative agarose gel images showing PCR amplification products are presented. (C) qRT-PCR analysis of transcript levels of the corresponding candidate genes in the T-DNA insertion lines. Three independent biological replicates were analyzed for genotype (n = 3). Each dot represents one independent biological replicate.

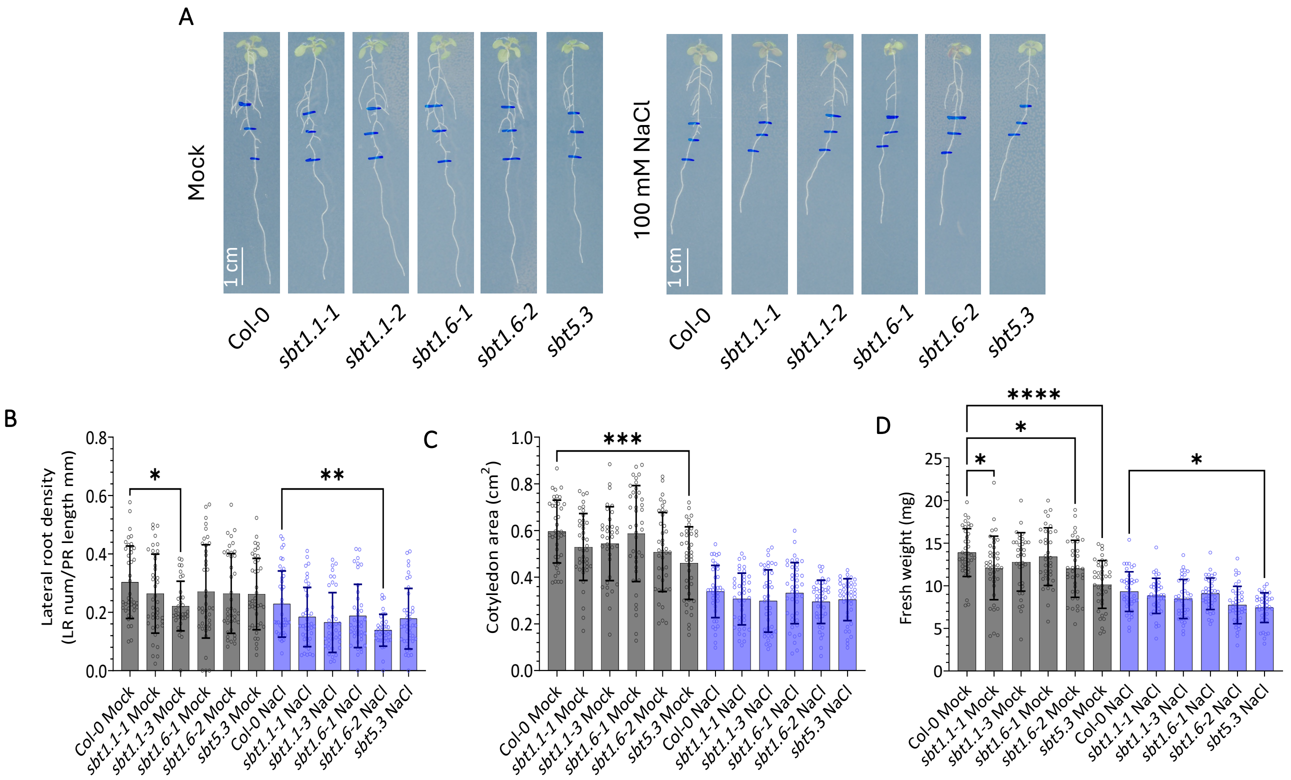


**Figure S4. Phenotypic analysis of *sbt* mutant seedlings under salt stress.** (A) Five-day-old seedlings grown on MS/2 medium supplemented with 1% Daishin agar and 0.5% (w/v) sucrose were transferred to control plates or plates supplemented with 100 mM NaCl. Primary root length was measured 5 d after transfer. (B) Lateral root density was determined 5 d after transfer. These seedlings correspond to those analyzed for lateral root number in Figure 2C. (C) Cotyledon area and (D) seedling fresh weight were measured 7 d after transfer. Data were obtained from two independent experiments, each including 20 seedlings per genotype and treatment (n = 40 seedlings in total per genotype and treatment). Each dot represents an individual seedling. Data are presented as mean ± SD. Statistical significance was assessed using one-way ANOVA followed by Šídák's multiple-comparisons test, comparing each mutant line with Col-0 within each treatment. Significance levels are indicated as * ($P < 0.05$), ** ($P < 0.01$), *** ($P < 0.001$), and **** ($P < 0.0001$).

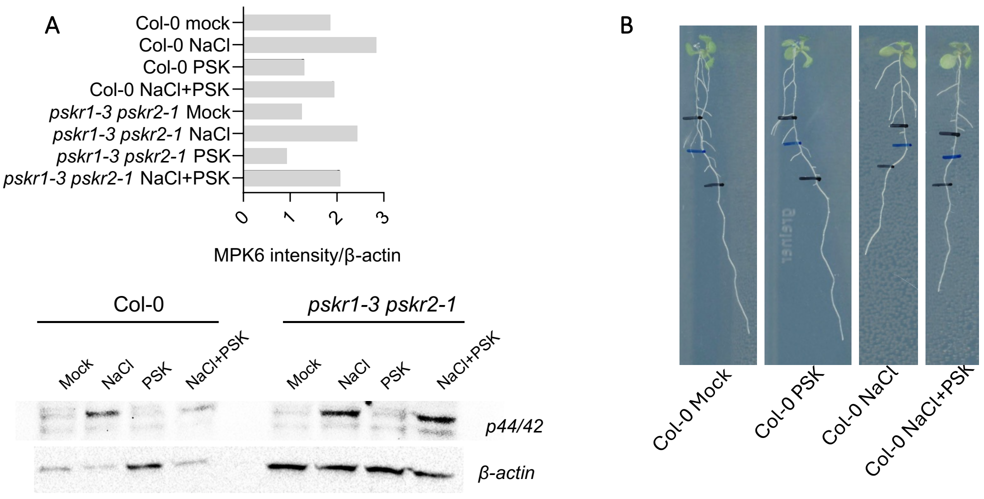


**Figure S5. Quantification of MPK6 phosphorylation and representative images of root growth under salt stress and PSK-α treatment.** (A) Quantification of MPK6 phosphorylation from the immunoblot shown in Figure 3B, together with an immunoblot from an independent biological replicate. Six-day-old Col-0 (wt) and *pskr1-3 pskr2-1* seedlings were treated for 15 min with mock medium or medium supplemented with 100 mM NaCl, 1 µM PSK-α or a combination of 100 mM NaCl and 1 µM PSK-α. MPK6 phosphorylation was detected by immunoblotting using an anti-p44/42 MAPK antibody. Band intensities were quantified and normalized to β-actin. (B) Representative images of Col-0 seedlings used for the root growth analysis shown in Figure 3E. Five-day-old seedlings were transferred to control plates or plates supplemented with 1 µM PSK-α, 100 mM NaCl, or a combination of 100 mM NaCl and 1 µM PSK-α and grown for an additional 5 d.

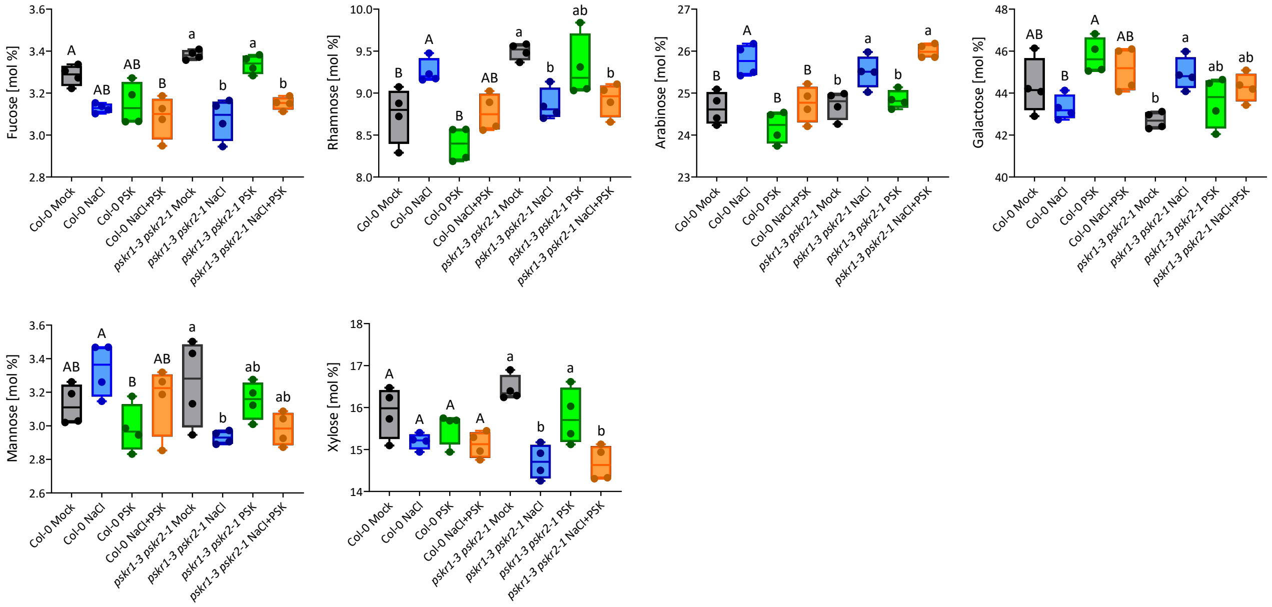


**Figure S6. PSK-α attenuates NaCl-induced changes in relative cell wall monosaccharide composition.** Six-day-old Col-0 and *pskr1-3 pskr2-1* seedlings grown in liquid MS/2 medium were treated for 48 h with mock medium or medium supplemented with 100 mM NaCl, 1 µM PSK-α, or a combination of 100 mM NaCl and 1 µM PSK-α. The relative neutral monosaccharide composition of cell wall fractions was determined by quantifying fucose, rhamnose, arabinose, galactose, mannose, glucose, and xylose, and expressed as mol% of total neutral sugars. Four independent biological replicates were analyzed for each genotype and treatment, with each replicate consisting of 40 seedlings ($n = 4$). Each dot represents an independent biological replicate. Data are presented as box-and-whisker plots showing the median and interquartile range ($25^{th}$–$75^{th}$ percentiles), with whiskers extending from the minimum to the maximum value. Statistical significance was assessed using one-way ANOVA followed by Tukey's HSD test to compare treatments within each genotype ($P < 0.05$). Different uppercase letters indicate significant differences among treatments in Col-0, whereas different lowercase letters indicate significant differences among treatments in *pskr1-3 pskr2-1*.
